# Nanoscale ligand density modulates gap junction intercellular communication of cell condensates during chondrogenesis

**DOI:** 10.1101/2021.04.28.441739

**Authors:** Ignasi Casanellas, Anna Lagunas, Yolanda Vida, Ezequiel Pérez-Inestrosa, Cristina Rodríguez-Pereira, Joana Magalhaes, José A. Andrades, José Becerra, Josep Samitier

## Abstract

To unveil the influence of cell-matrix adhesions in the establishment of gap junction intercellular communication (GJIC) during cell condensation in chondrogenesis.

**Materials & Methods:** Previously developed nanopatterns of the cell adhesive ligand arginine-glycine-aspartic acid (RGD) were used as cell culture substrates to control cell adhesion at the nanoscale. We conducted *in vitro* chondrogenesis of mesenchymal stem cells on the nanopatterns. We evaluated cohesion and GJIC in cell condensates.

**Results:** Mechanical stability and GJIC are enhanced by a nanopattern configuration in which 90% of the surface area presents adhesion sites separated less than 70 nm, thus providing an onset for cell signaling.

**Conclusion:** Cell-matrix adhesions regulate GJIC of mesenchymal cell condensates during *in vitro* chondrogenesis from a threshold configuration at the nanoscale.

## Introduction

Mesenchymal cell condensation is a critical morphogenetic transition regulated by cell adhesion in which mesenchymal stem cells (MSCs) gather to form intimate cell-to-cell contacts [1]. The condensation phase was named “membranous skeleton” by Grüneberg to stress its distinctive function in skeletal development [2]. During skeletogenesis, cell density increases locally at the condensation sites by means of extracellular matrix (ECM)-driven active cell movement [1], with an important role of the ECM ubiquitous protein fibronectin (FN), which is upregulated during mesenchymal cell condensation [3,4].

In osteochondral development, mesenchymal cell condensation is concurrent with the formation of an extensive gap junction (GJ) communication network [5]. GJs are plasma membrane channels that provide intercellular communication in almost all animal tissues, allowing cells to exchange ions and small molecules through a controlled gating mechanism [6]. During morphogenesis, an efficient network of GJs is an extremely versatile communication system that mediates the rapid synchronization between cells. GJs established during embryonic patterning allow multicellular groups to coordinate towards the formation of supracellular, tissue-level structures. Avascular tissues, such as cartilage, particularly rely on this form of intercellular communication for successful development [7–9]. Connexin 43 (Cx43) is the most widely expressed and studied GJ protein [10] and modulates cartilage structure through its C-terminal cytosolic domain [11].

Although previous studies have related gap junction intercellular communication (GJIC) with matrix-associated proteins [12–15], little is known about how extracellular inputs modulate the formation of intercellular GJ networks and the associated implications in tissue architecture and function. Integrin-mediated cell-matrix interactions regulate many biological processes such as cell shape, proliferation, migration, differentiation and programmed cell death [16–18]. During morphogenesis, dynamic adhesion mechanisms, together with the associated regulatory signalling pathways, define tissue differentiation and architecture, and modulate collective cell behaviour [19–21].

Cell adhesion is governed at the nanoscale, as evidenced by the assembly of collagen fibres in the ECM or the folding/unfolding of fibronectin [22–24], and also by the compartmentalization of the cell membrane receptors into nanoclusters, which facilitates allosteric regulation, increases the ligand rebinding probability and triggers the assembly of signalling complexes in the cytosol [25]. Studies on cell culture platforms nanopatterned with ECM motifs have shown the importance of ligand presentation in modulating cell-substrate interactions at the nanoscale. Micellar lithography was employed to produce nanopatterns of the cell-adhesive peptide arginine-glycine-aspartic acid (RGD), present in fibronectin, which revealed a nanospacing threshold of around 70 nm for an efficient cell adhesion on rigid substrates and demonstrated that local ligand presentation affects cell adhesion more than the global ligand density [26–29]. More recent works showed that not only cell adhesion, but also many other cell responses are determined by the nanoscale ligand presentation [30].

We have previously described a nanopatterning technique for cell adhesion studies in which RGD-functionalized dendrimers were adsorbed on Poly(L-lactide) (PLLA) to generate uneven nanopatterns with liquid-like order and defined spacing on large areas, thus being fully compatible with standard cell culture protocols [31]. Owing to steric hindrance restrictions, each dendrimer of 4-5 nm in diameter, although bearing up to eight copies of the RGD ligand, provides a single site for integrin receptor binding. Therefore, dendrimer nanopattern configuration directly correlates with the RGD available for cell adhesion. By varying the initial concentration of dendrimers in solution, different nanopattern configurations can be obtained, with different degrees of substrate cell adhesiveness. These nanopatterns sustain cell adhesion more efficiently than the corresponding homogeneous surfaces [31] and improve cell differentiation [32–34].

Taking chondrogenesis as a model [35], we used the RGD dendrimer-based nanopatterns to conduct a systematic study a systematic study on the influence of the nanoscale surface adhesiveness in the establishment of GJIC during mesenchymal cell condensation in *in vitro* chondrogenesis. Results show that mesenchymal cell condensation produced under chondrogenic stimuli depends on integrin-based cell-matrix adhesions, and that the mechanical stability and GJIC of cell condensates are enhanced by a nanopattern configuration in which 90% of the surface area presents adhesion sites separated less than 70 nm, thus providing a threshold for cell signalling. Substrate information is continuously propagated within cell condensates through cytoskeletal contractility.

## Materials & methods

### Production of nanopatterned substrates

All steps were performed in a sterile tissue culture hood, and only sterile materials, solutions and techniques were used. Nanopatterned substrates were prepared as previously described [32,33]. Corning^®^ glass microscopy slides of 7.5 x 2.5 cm (Sigma-Aldrich, 2947-75X25) were cut with a diamond-tip cutter to square pieces of 1.25 x 1.25 cm. Slides were washed thoroughly with deionized water (18 MΩ·cm Milli-Q, Millipore) followed by 96% ethanol and air-dried. Glass substrates were brought to 60°C for 10 min and spin-coated with a poly(L-lactic acid) (PLLA) (Corbion) 2% m v^-1^ solution in dry 1,4-dioxane (Sigma-Aldrich, 296309). Slides were coated with a two-step program: 5 s at 500 rpm with an acceleration of 300 rpm/s (to eliminate excess solution and spread the remaining solution homogeneously on the surface) followed by 30 s at 3000 rpm with an acceleration of 1500 rpm.

Dendrimer solutions were prepared in deionized water at 2.5×10^-8^, 10^-8^ and 4×10^-9^% w w^-1^ concentrations, sonicated for 10 min and filtered through a 0.22 μm Millex RB sterile syringe filter (Merck Millipore) attached to a syringe. Glass substrates were sterilized in UV light and then immersed in dendrimer solutions for 16 h (pH = 5.6, T = 293 K). Substrates were then rinsed with deionized water and transferred to a new culture plate. FN-coated substrates (S_FN_) were produced by incubating PLLA substrates in FN from bovine plasma solution (Sigma-Aldrich, F1141) 100 μg/ml phosphate buffered saline (PBS, Gibco, 21600-10) for 1 h at room temperature, just before cell seeding.

### Cell culture

As previously described [36], human adipose-derived mesenchymal stem cells (hASCs) (ATCC, PCS-500-011) were cultured in MSC Basal Medium (ATCC, PCS-500-030) supplemented with MSC Growth Kit Low Serum (ATCC, PCS-500-040). Medium was replaced every 2 days. Passaging was carried out when cells reached 70-80% confluence. For the experiments, cells were trypsinized at passages 3 to 4, counted, resuspended in chondrogenesis-inducing medium Chondrocyte Differentiation Tool (ATCC, PCS-500-051) with 0.1% v v^-1^ penicillin-streptomycin (Invitrogen, 15140), and seeded on nanopatterned and control substrates at a density of 2,760 cells cm^-2^. 3 replicates of each condition were seeded. Medium was replaced every 3 days.

### Immunostaining

After 6 and 9 days of chondrogenic differentiation on the nanopatterns and controls, cells were carefully rinsed with PBS (Gibco, 21600-10) and fixed with Formalin Solution (Sigma, HT5011) for 20 min at room temperature. Samples were blocked with 50 mM ammonium chloride (Sigma, A9434) in PBS for 20 min and permeabilized with saponin (Sigma, 47036) 0.1% m v^-1^ in Blocking Solution (BSA (Sigma, A3059) 1% m/v in PBS) for 10 min. Samples were stained with primary antibodies against VCL (Abcam, ab18058) or Cx43 (Abcam, ab63851) at 5 μg mL-1 in Blocking Solution for 1 h at room temperature, then washed with PBS and treated with fluorophore-conjugated secondary antibodies (according to the organism of the primary antibody): anti-rabbit Alexa 568 (LifeTech, A11036) or anti-mouse Alexa 488 (LifeTech, A10667) 1 μg/mL in Blocking Solution for 1 h at room temperature, covered from light. For nuclei observation, Hoechst 33342 (Invitrogen, H3570) 1:1000 was included in the secondary antibody solution. Samples were washed with PBS and left to dry on air for 5-10 min, then placed on a Corning® 7.5 x 2.5 cm microscopy slide. One drop (50 μl) of Fluoromount mounting medium (Sigma, HT5011) was added, and a Corning® 1.8 x 1.8 cm coverslip (Sigma-Aldrich, 2845-18) was placed on the sample avoiding the formation of bubbles. Mounted samples were stored at 4°C covered from light. At least 16 h were left to elapse before sample imaging.

### Image acquisition and analysis

Samples were imaged with a Leica SPE Upright Confocal Microscope (Leica Microsystems) with a 40x/1.15 NA objective. The distance between imaged slices (z-size) was set at 1 μm. At least 3 cell condensates were imaged for each sample except on fibronectin-coated substrates, where cells developed a monolayer and did not always generate 3 condensates.

Images were analyzed with ImageJ software. For condensate size measurements, a z-projection of each condensate was created, and the whole condensate area was manually selected and measured. Confocal images stained for Hoechst at day 6 were used to measure internuclear distances between adjacent cells in condensates. Slides in the central region of condensates were selected for analysis and the straight-line horizontal distances between the centers of adjacent cells were measured.

For the measurement of Cx43 expression confocal z-projections were used (maximum stained area per sample). The background of z-projections was removed, and a threshold was applied to select areas of Cx43 expression. The obtained total area was normalized against the area of the corresponding condensate.

To analyze Cx43 network connectivity, a threshold was applied to the confocal stacks of Cx43 immunostaining, to obtain 3D binary images (stacks of 8-bit images). Binary stacks were then processed with the Skeletonize plugin in Fiji (Fig. S1, Video S1) to generate the protein network’s skeleton. The skeleton of a shape (in this case, protein staining) is defined as a thin representation that is equidistant to the shape’s boundaries, and which contains all the information necessary to reconstruct the shape (such as connectivity, topology, length, or direction). We analyzed the resulting Cx43 network skeletons with the Analyze Skeleton plugin in Fiji, which tags all pixel/voxels in the skeleton image and then counts all its junctions, triple and quadruple points and branches, and measures their average and maximum length. We retrieved the number of end-point voxels (pixels with less than two neighbors) and the mean branch length in each condensate and normalized them to the Cx43 immunostained area and the number of slices taken for analysis in the initial stack. Network connectivity was calculated as the inverse value of the end-point voxels per volume unit of Cx43 staining.

### RNA extraction and retrotranscription

Reverse transcription real-time PCR (RT-qPCR) was performed to measure *VCL* and Cx43 (*GJA1*) expression at 6 and 9 days of chondrogenesis. All working surfaces and tools were treated with RNAse Zap decontamination solution (Thermo Fisher Scientific, AM9780) prior to any steps involving RNA. Three cell culture replicates of each condition were seeded as described above. Messenger RNA (mRNA) was extracted from the samples and purified with a RNeasy Micro Kit (Qiagen, 74004), then quantified in a Nanodrop ND-1000 Spectrophotometer (Thermo Fisher Scientific). mRNA was retrotranscribed into cDNA with an iScript Advanced cDNA Synthesis Kit (Bio-Rad, 1725037) in a T100 Thermal Cycler (Bio-Rad). The same procedure was performed on undifferentiated human mesenchymal stem cells (hASCs) as a reference.

### qPCR and data analysis

qPCR was performed as previously described [36], with the Sso Advanced Universal SYBR Green Supermix kit (Bio-Rad, 1725271) in an Applied Biosystems StepOnePlus Real-Time PCR Machine (Thermo Fisher Scientific). Commercial primer pairs were used for *VCL* (Bio-Rad, qHsaCID0020885) and *GJA1* (Bio-Rad, qHsaCID0012977), as well as *B2M* (Bio-Rad, qHsaCID0015347) and *RPL24* (Bio-Rad, qHsaCID0038677) as housekeeping genes. To prevent gDNA amplification, a DNase digestion step was included during RNA extraction and intron-spanning primer pairs were selected. The amplification program consisted of an initial activation step of 30 s at 95°C, followed by 50 cycles of 10 s at 95°C for denaturation and 1 min at 60°C for annealing and extension, and a final denaturation step of 15 s at 95°C. Melt curves were performed from 65°C to 95°C in steps of 0.5°C. Technical duplicates of each sample were performed in the qPCR.

qPCR data were analyzed with qBase+ software version 3.1 (Biogazelle, Zwijnaarde, Belgium). Only samples with a Ct below 40 were considered for analysis. The expression of each gene was calculated by the 2^-ΔΔCt^ method, normalized to that of undifferentiated hASCs (assigned value 1) and presented as relative mRNA expression levels.

### Western blotting

At day 6 of chondrogenesis, samples were rinsed with PBS and cells were lysed with RIPA buffer (150 mM NaCl, 5 mM EDTA, 50 mM Tris-HCl pH=8, 1% Triton X-100, 0.1% SDS, EDTA-free Protease Inhibitor Cocktail) on ice for 45 min. Cell lysate solutions were centrifuged at 16100 g for 10 min at 4°C and the pellet was discarded. Total protein concentration was quantified with a Pierce TM BCA Protein Assay Kit (ThermoFisher, 23227) in a Benchmark Plus Microplate Reader (Bio-Rad).

Protein samples (5 μg, volume calculated according to each sample’s total protein concentration) were mixed with loading buffer. The volume was adjusted to 20 μl with MQ water, samples were briefly vortexed and spinned and heated to 96°C for 10 min. Samples were loaded in each well of an SDS-PAGE gel (Mini-Protean TGX Precast Gel 12%, Bio-Rad, 456-1045) along with a protein weight ladder. Protein electrophoresis was performed in a vertical cell (Mini-Protean System, Bio-Rad). Voltage was set at 50 V until samples transitioned from the stacking to the running section of the gel, then at 60 V for 1.5-2 h.

The resulting gels were transferred to PVDF membranes (Amersham Hybond, 10600023) for 2 h at 60 V at 4°C. After the transfer, membranes were stained with Ponceau solution; images were taken to quantify total protein amount in each lane.

For immunostaining, membranes were blocked with dry milk dissolved in Tris Buffer Saline with 1% Tween 20 (TBST) for 1 h and probed overnight at 4°C with mouse anti-VCL (Abcam, ab18058) and rabbit anti-Cx43 (Abcam, ab63851) primary antibodies, followed by IgG HRP-linked secondary antibodies anti-mouse (Cell Signaling, 7076) and anti-rabbit (Cell Signaling, 7074) in 3% BSA in TBST for 1 h at room temperature. Immunoblots were developed using Clarity ECL Western substrate (Bio-Rad, 1705060). Bands were visualized in an ImageQuant LAS 4000 imager (GE Healthcare). Integrated density of bands was measured with Fiji. Background signal was measured in empty areas of the blot and subtracted from the corresponding values. Protein production in each lane was normalized to the integrated density of total protein staining.

### Neurobiotin (NB) assay

A tracer assay was performed to analyze the functionality of GJIC networks. MSCs were seeded in chondrogenesis-inducing medium, as described above. After 6 days of differentiation, samples were washed with HBSS buffer without calcium or magnesium (Life Technologies, 14175095) and treated with neurobiotin 2% m v^-1^ (Vector, SP-1120) in HBSS for 90 s at 37°C. Samples were then washed with HBSS, fixed with Formalin Solution, permeabilized with saponin and stained with Streptavidin-Texas Red conjugate (Life Technologies, S872) and Hoechst 1 μg mL^-1^ in BSA 1% m v^-1^ in PBS for 1 h at room temperature. Samples were imaged with a Leica SPE Upright Confocal Microscope as described above.

Images were analyzed with ImageJ software. A z-projection of each condensate was created, and background was removed. Distance of neurobiotin spread was measured in a straight line from the outer rim of the condensates inwards, in at least two separate locations for each condensate.

### Fluorescence recovery after photobleaching (FRAP) microscopy

hASCs were cultured in chondrogenic medium as described on S_18_ or S_90_ nanopatterns. To block GJ accretion, 18β-Glycyrrhetinic acid (Sigma, G10105) 50 μM was added on S_90_ samples with a change of medium, 15 h before tracer loading. At least two samples of each condition were seeded and analyzed.

At day 6 of differentiation, samples were incubated for 5 minutes in HBSS. Fluorescent neurobiotin-488 (Vector, SP-1125-2) was dissolved in HBSS to 1 mg/ml, added to the samples and left to incubate at 37°C for 20 min. Samples were then washed twice with HBSS and kept in chondrogenic medium at 37°C until imaging.

Samples were imaged immediately after tracer loading at 37°C and 5% CO_2_, in a Leica LSM780 confocal microscope with a 63X objective, in glass-bottom microscopy dishes (VWR, 734-2906). The pinhole was kept open to the maximum (11 Airy Units). Images were taken every 15 seconds for a total of 10 minutes. Five images were taken before bleaching. Bleaching was then performed with 5 iterations of 488 laser at full power, with a pixel dwell time of 100 μs. Three regions of interest (ROIs) were followed for analysis: the bleaching region (consisting of a circle of 30 pixels of diameter in a central area of a condensate), the region of total fluorescence (normally taking the whole cell condensate, where available), and a background region outside the condensate. FRAP data analysis was performed with the online tool EasyFRAP (https://easyfrap.vmnet.upatras.gr/) [37]. Intensity results were normalized with the Full-Scale method to values between 0 and 1 and graphed over time as percentages of the pre-bleaching fluorescence. Quantitative analysis was conducted by linear regression of the fluorescence recovery values to an exponential decay function as previously described [38–40].

### Condensate transplantation assay

Nanopatterns with 18% (S_18_) and 90% (S_90_) of local surface adhesiveness were used (Fig. S2 and Table S1). A transplantation assay was performed to study the effects of RGD nanopatterned substrates on formed condensates and the propagation of the adhesive information from the substrate into cell condensates. Cells were cultured on the nanopatterns in chondrogenesis-inducing medium as described above. After 3 days, cell condensates formed on the nanopatterns of S_90_ were removed by pipetting and transferred to new S_90_ or S_18_ substrates. Transplanted condensates were cultured on the new substrates for another 3 days, to a total of 6 days of differentiation. For each sample, around half of the condensates were transplanted, whereas the other half were kept on the original substrate (not transplanted) as a control of unaltered differentiation. Three replicates of each condition were seeded and transplanted.

Samples were fixed, immunostained, imaged and analyzed for Cx43 expression as described above. Results were normalized to those of non-transplanted S_90_ condensates (assigned value 1) and presented as relative values.

To visualize condensates from the side (transversal cuts), Z-stacks were resliced and one image from the center of the condensate was selected. Cx43 production at the basal and top layers of condensates was measured by the average staining intensity at each region on unprocessed images. For each condensate, average basal intensity was divided by average apical intensity to obtain the Basal/Apical ratio, indicative of Cx43 distribution within condensates.

### Integrin blocking and myosin inhibition assays

Three replicates of each condition were seeded in chondrogenesis-inducing medium as described above. For integrin blocking samples, medium was changed to fresh medium containing RGD dendrimers in solution at 4 10^-9^% w w^-1^ after 5 days of differentiation, 24 h before fixation. We selected this dendrimer concentration because it yields S_18_ substrates. During substrate functionalization, equilibrium is reached between dendrimer concentration in solution and adsorbed dendrimer density; hence, use of the concentration corresponding to the substrates with the lowest density prevents further adsorption mid-assay.

For the myosin inhibition experiment, medium was changed to fresh medium with 50 μg mL^-1^ blebbistatin (Sigma, B0560) 6 h before fixation.

All samples were fixed at day 6 of chondrogenesis, immunostained with anti-Cx43 antibody and Sir-Actin (Tebu-bio, SC001), and imaged with a Zeiss LSM780 Confocal Microscope (Zeiss Microscopy) with a 40x objective. Cx43 expression in condensates was quantified as described above and normalized to corresponding non-treated samples.

### Statistics

Quantitative data are displayed, showing average and standard error of the mean (SEM). n is the sample size. At least 3 independent substrates were analyzed for each condition. Significant differences were judged using the One-way ANOVA with Fisher LSD post-hoc test or T-test when only two groups are compared, using OriginPro 8.5. Where data did not pass a normality test, a Kruskal-Wallis test with Dunn means comparison was applied with GraphPad Prism 8.3. A *p* of less than 0.05 was considered statistically significant.

## Results

### Measurement of cell-cell cohesion in mesenchymal cell condensates

We used three different nanopattern configurations with increasing adhesiveness: S_18_, S_45_ and S_90_ (subindexes indicate the percentage of surface area containing adhesion sites separated less than 70 nm; Fig. S2 and Table S1). Pristine non-patterned PLLA (S_0_) and FN-coated PLLA (S_FN_) substrates were the negative and positive controls for cell adhesion, respectively [32]. hASCs were seeded on the nanopatterns in chondrogenic medium. To evaluate cell-cell cohesion in cell condensates, we measured the distance between the cell nuclei in adjacent cells, and the expression of the adhesome protein vinculin (VCL) at day 6 of chondrogenic induction. VCL is an actin-binding cytoskeletal protein present in cell-matrix and cell-cell adhesions. It stabilizes the dynamics of adherens junctions (AJs) by binding to actin filaments, thereby reinforcing cell-cell interactions [41]. Moreover, VCL is known to stabilize Cx43-containing GJs in cardiac myocytes through binding to the scaffolding protein zonula occludens-1, and thereby acting as a binder between adherens junctions and GJs [14]. We found that condensates on S_90_ and S_FN_ were spatially distributed closer together than those on S_0_, S_18_ and S_45_, indicating that condensates on high-adherence substrates were more tightly packed (Fig. 1a). *VCL* expression showed a tendency to increase with decreasing substrate adhesiveness (Fig. 1b). At days 6 and 9 of chondrogenic induction, *VCL* was upregulated on S_0_ and S_18_ substrates with respect to S_FN_, where the expression levels were comparable to those of the undifferentiated hASCs. Western blotting of VCL at day 6 of chondrogenic induction showed increased production of the protein with decreasing surface adhesiveness (Fig. 1c and S3a), thus mirroring mRNA expression results. Immunofluorescence images (Fig. 1d) showed that VCL accumulated at the rim of cell condensates, wrapping the cells in a compacted structure.

**Figure 1.**
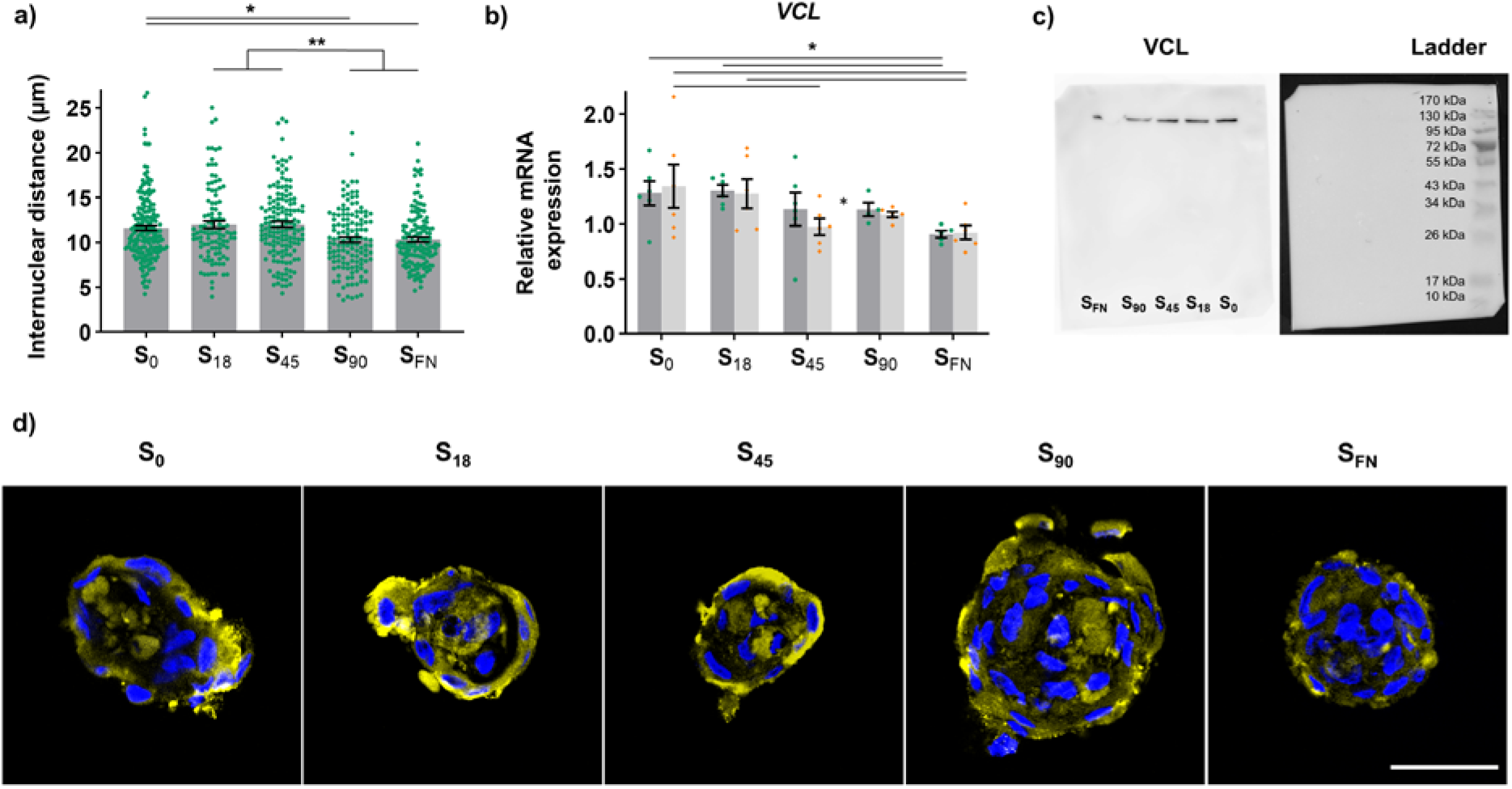
Cell cohesion in the mesenchymal cell condensates obtained on the different substrates was evaluated by **a)** measuring the distance between cell nuclei in adjacent cells in cell condensates at day 6 of chondrogenic induction (106 ≤ n ≤ 192), and **b)** through the expression of vinculin mRNA (*VCL*), relative to that of undifferentiated hASCs (assigned value 1, not shown) at day 6 (dark bars with green dots) and day 9 (light bars with orange rhomboids) of chondrogenic induction (n = 6). Results are given as the mean ± SEM, *p < 0.05, **p < 0.01. **c)** Western blot results for VCL obtained at day 6 of chondrogenic induction. **d)** Representative confocal microscopy images of condensates stained for VCL (yellow) and Hoechst (blue) at day 6 of chondrogenic induction. Scale bar = 40 μm.

### Connexin-43-based gap junction formation

GJs are formed by two opposed connexons or hemichannels docked end-to-end in adjacent cells. Each connexon is composed by 6 connexin (Cx) subunits (Fig. 2a). To examine cell-cell interconnectivity through GJ formation in the cell condensates, we first measured Cx43 expression. We analysed the expression of *GJA1* (the gene encoding for Cx43) at day 6 and 9 of chondrogenic induction (Fig. 2b). *GJA1* was overexpressed on S_90_ to over twice the level of undifferentiated hASCs at day 6. However, expression on S_90_ levelled off at day 9 among the different nanopattern configurations and S_0_. *GJA1* expression in S_18_ and S_45_ nanopatterns was comparable to that of the negative control both at day 6 and 9 of chondrogenic induction. GJA1 was downregulated on S_FN_.

**Figure 2.**
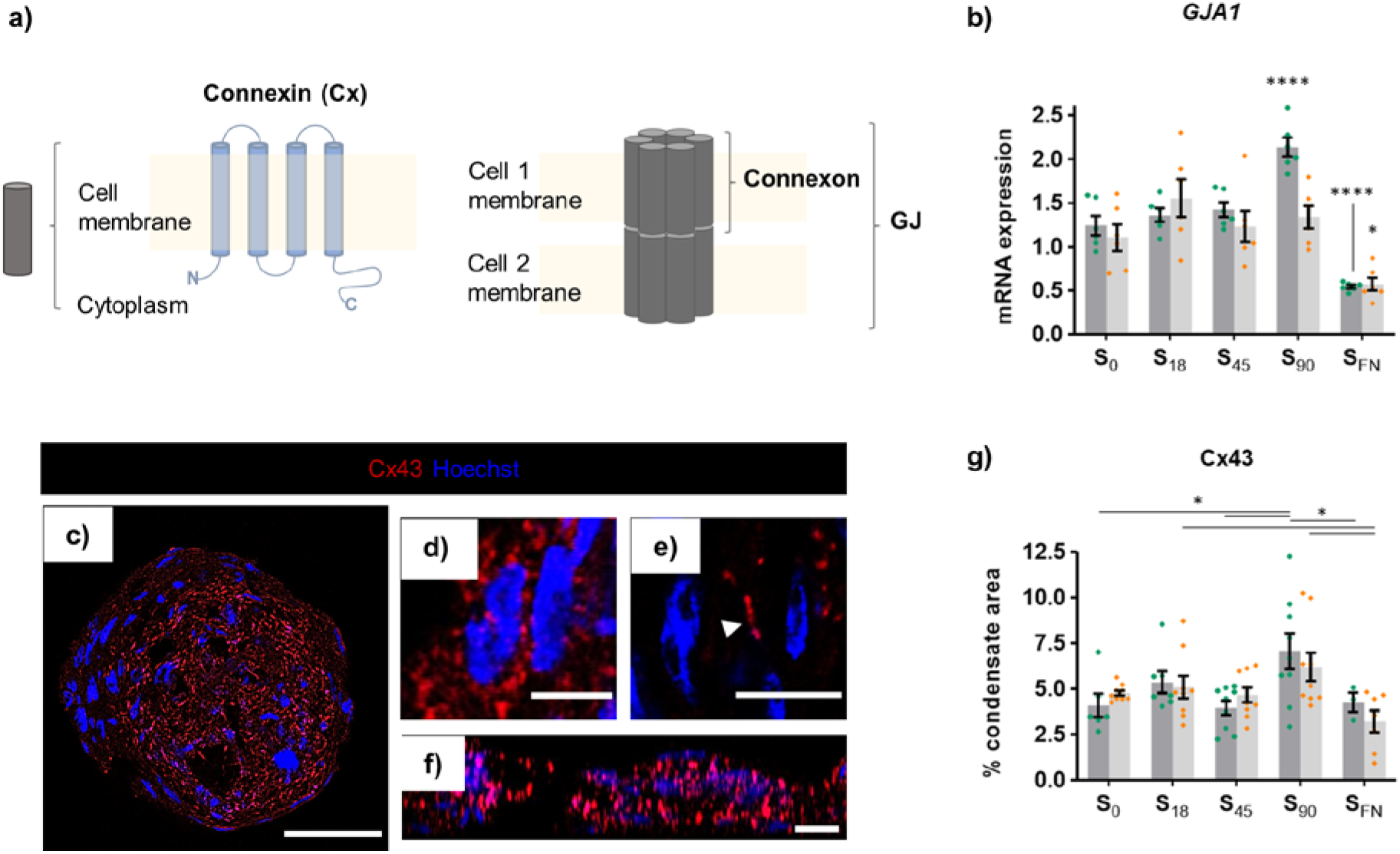
Cx43 expression in the cell condensates. **a)** Schematic representation of GJ structure. **b)** Expression of *GJA1* mRNA, relative to that of undifferentiated hASCs (assigned value 1, not shown) (n = 6) and **c)** Representative confocal slice taken at the mid-height of a cell condensate on S_45_ nanopattern. Cell nuclei are in blue (Hoechst staining) and Cx43 in red, showing the formation of Cx43 aggregates of different sizes and locations in cells within the cell condensates. Scale bar = 50 μm. **d)** Round aggregates of around 1-2 μm in diameter located at the rim of cell condensates. **e)** Larger aggregates of up to 5-10 μm located at the center of cell condensates, at the interfaces between cells. Also visible in the YZ projection in **f).** Scale bars = 10 μm. **g)** Quantification of Cx43 production from immunostaining in cell condensates (3 ≤ n ≤ 9) at day 6 (dark bars with green dots) and day 9 (light bars with orange rhomboids) of chondrogenic induction. Results are given as individual sample values with the mean ± SEM, *p < 0.05, ****p < 0.0001.

We also immunostained samples for Cx43 (Fig 2c-f). We observed Cx43 aggregates of different sizes are formed at different locations in cells within the mesenchymal cell condensates (Fig. 2c). Round aggregates of around 1-2 μm in diameter could be found mostly at the rim of cell condensates (Fig. 2d), while larger aggregates of up to 5-10 μm locate preferentially at the center of cell condensates, at the interfaces between cells (Fig. 2e and f). Quantification of the percentage of immunostained area for Cx43 in the cell condensates (Fig. 2c) from confocal z-projections (Fig. S3) at day 6 of chondrogenic induction showed higher values on S_90_ when compared to most other substrates. A slight decrease in the percentage of immunostained area in S_90_ cell condensates was observed at day 9 of chondrogenic induction, in agreement with mRNA expression results (Fig. 2b). We examined total Cx43 production by Western blot after 6 days of chondrogenic induction (Fig. S4). We found that protein production increased with nanopattern adhesiveness. Surprisingly, a high protein content was detected also for S_FN_ controls, in contrast to the relative mRNA expression, which was significantly lower for S_FN_ than for the rest of the substrates at day 6 (Fig. 2b). This was attributed to differences in the configuration adopted by cells in S_FN_ substrates, where they are mostly in a monolayer configuration compared to the cell condensates found on nanopatterns and S_0_, thereby making the extraction process particularly more efficient for Cx43 in S_FN_ substrates and causing a bias in the Western blot result.

Connexin hemichannels, or connexons, accumulate and dock with apposed connexons from adjacent cells forming dense GJ plaques. GJIC has been found to largely depend on the size of these plaques [42], which are continuously regenerated by the addition of connexon subunits at the edges and internalization from the center of the plaques [7,43,44]. Such a continuous renewal may ensure the maintenance of the established GJIC network. Skeletonization of Cx43 immunostained images renders a 3D representation of the distribution of the protein forming clusters, either GJs or arrayed connexons (Fig. 3a, Fig. S1 and Video S1), from which the average branch length (Fig. S5) and number of branch terminations (end-point voxels; EPVs) can be calculated. Shorter branches and fewer end-point voxels indicate a more intricately shaped network, as shown in the zoomed-in sections of the skeletonized images of cell condensates from S_90_ and S_0_ nanopatterns (Fig. 3a). Network connectivity (calculated as the inverse of EPVs number per protein volume unit) showed a tendency to increase with substrate adhesiveness up to S_90_ and then decrease for S_FN_ controls. At day 9 of chondrogenic induction, connectivity in S_90_ cell condensates was significantly higher than in S_0_, S_18_ and S_FN_.

**Figure 3.**
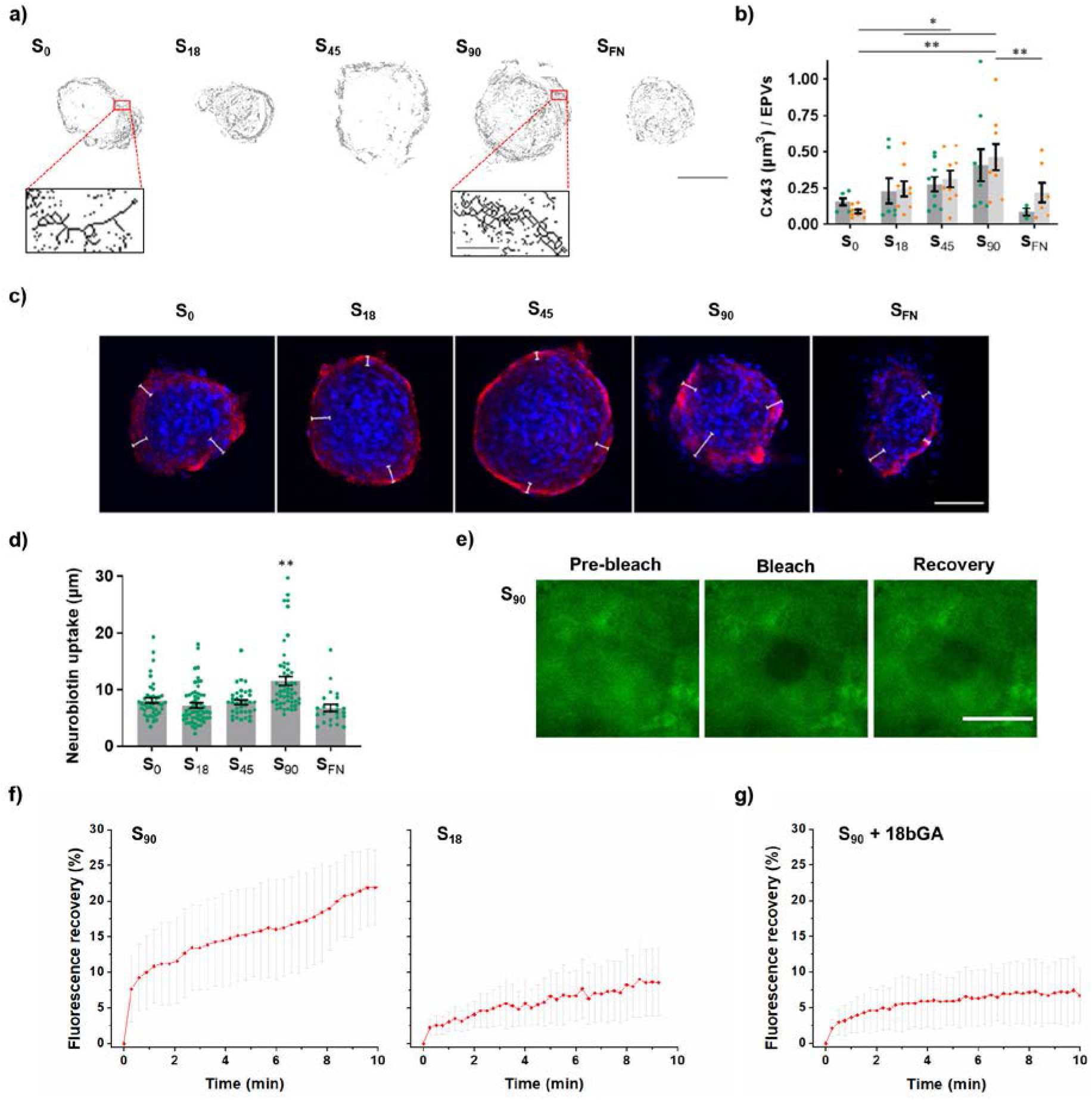
**a)** Top: Skeletonized z-projections of Cx43 immunostaining; scale bar = 40 μm. Bottom: zoomed-in sections of S_0_ and S_90_; scale bar = 3 μm. **b)** Network connectivity calculated as the inverse values of the number of end-point voxels (EPVs) normalized to the volume of Cx43 staining in confocal images (3 ≤ n ≤ 9) at day 6 (dark bars with green dots) and day 9 (light bars with orange rhomboids) of chondrogenic induction. **c)** Confocal z-projections showing neurobiotin tracer (red) and cell nuclei (blue) after 90 s of exposure at day 6 of chondrogenic induction. White lines are the measured distances of tracer uptake. Scale bar = 80 μm. **d)** Quantification of tracer uptake (24 ≤ n ≤ 58) after 90 s exposure. Results given as the mean ± SEM, *p < 0.05, **p < 0.01. **e)** Representative fluorescence images of FRAP microscopy conducted on S_90_ cell condensates. Scale bar = 20 μm. **f)** Percentage of fluorescence recovery of S_90_ and S_18_ cell condensates after photobleaching. (43 ≤ n ≤ 60). **g)** To confirm the involvement GJs in the intercellular transfer of NB, GJs coupling was blocked with 18βGA (S_90_ + 18βGA), leading to a 77 % decrease of fluorescence recovery.

Observation of the spread of biotinylated or fluorescent tracers has become one of the most common methods for demonstrating GJs network coupling [42][45] [46]. NB is a GJ/hemichannel permeable dye that can penetrate from the exposed Cx43 connexons and diffuse inwards across the GJ network when calcium ion concentration is maintained below the physiological levels (open channel conformation). To assess GJIC in the cell condensates on the different substrates, we conducted a NB tracer uptake assay. We first treated cell condensates of 6 days of chondrogenic induction with NB in a buffered solution in the absence of calcium for 10 minutes to allow the tracer permeation through the exposed hemichannels. However, this resulted in condensates on all conditions almost completely filled with tracer, which made any differences in the uptake distance hardly observable (Fig. S6). We then reduced the time of exposure to 90 seconds (Fig. 3c). Quantification of NB diffusion into cell condensates showed the uptake is significantly higher in S_90_ nanopatterns, while statistically equal on all other substrates (Fig. 3d). The mean uptake rate of NB in condensates ranged from 4.5 μm/min on S_FN_ to 7.7 μm/min on S_90_.

To directly measure GJIC in the cell condensates, we conducted fluorescence recovery after photobleaching (FRAP) microscopy [42,48]. Cell condensates from S_90_ and S_18_ nanopatterns of 6 days of chondrogenic induction were treated with fluorescently labelled NB for 20 minutes to ensure the preloading of the condensates with dye. A selected group of cells in cell condensates was photobleached with a brief pulse of high-intensity laser illumination, and the fluorescence recovery of the photobleached region was recorded to visualize the transfer of NB from adjacent cells (Fig. 3e and 3f). Quantitative assessment of FRAP was carried out by linear regression of fluorescence recovery values to an exponential decay function commonly used for analysis of gap-FRAP experiments [38–40]. We obtained an average fluorescence recovery of 30% for S_90_ cell condensates, and 11% for S_18_ cell condensates, with fluorescence half-recovery times of 5.3 and 4.8 min, respectively. To particularly address GJ-dependent exchange of NB, we pre-treated S_90_ cell condensates with 18β-glycyrrhetinic acid (18βGA), a saponin that induces disassembly of GJ plaques through Cx dephosphorylation [49,50]. We observed that the fluorescence recovery in this case was drastically reduced (Fig. 3g). Fitting of the fluorescence recovery data lead to a 7% with a T half of 1.9 min (77 % reduction compared to S_90_ without 18βGA pre-treatment), a similar value to that obtained for S_18_ cell condensates. This result indicates that GJs coupling is responsible of the higher values of fluorescence recovery obtained for S_90_, and that GJ coupling is very low or negligible in S_18_ cell condensates, where dye transfer most likely occurs by passive diffusion or mediated by another route.

### Integrin-mediated regulation of connexin-43

Results show that cell condensates on S_90_ nanopatterns developed a more efficient Cx43-based GJ network, thus suggesting integrin-based adhesions play a role in the development of GJIC during mesenchymal cell condensation in chondrogenesis. To specifically address the role of integrins, we added RGD-functionalized dendrimers in solution on formed condensates (5 days of chondrogenic induction) to block integrin receptors at the cell membrane. Disturbance of integrin clustering at cell-substrate adhesion sites led to a decrease in the percentage of area immunostained for Cx43 in S_90_ cell condensates but not in S_18_ (Fig. 4a). This demonstrates that the effects of S_90_ nanopattern configuration are mediated by integrins and that they are sustained at least for 5 days in culture.

**Figure 4.**
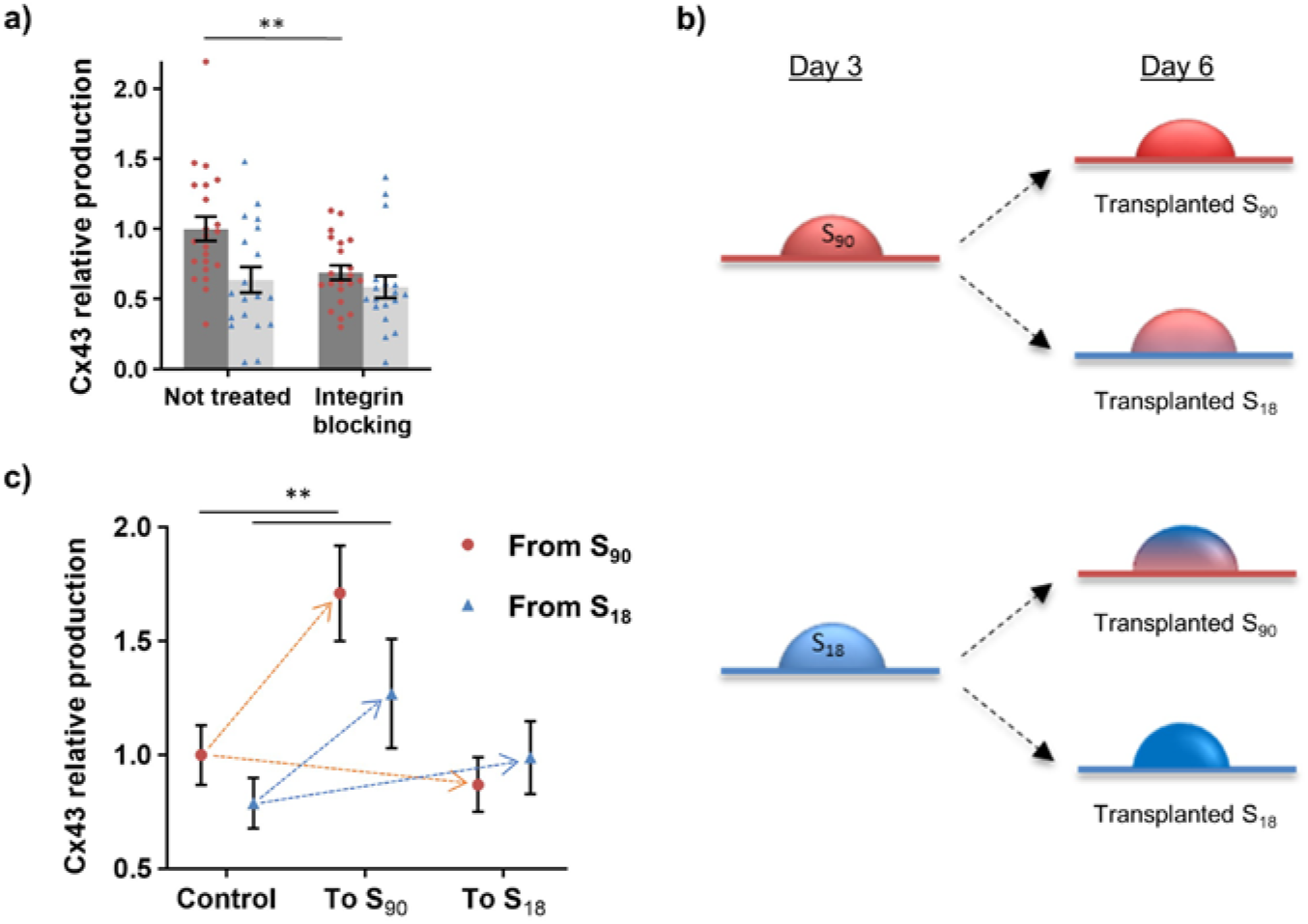
**a)** Quantification of the relative area immunostained for Cx43 on S_90_ (dark bars with red dots) and S_18_ (light bars with blue triangles) obtained for non-treated cell condensates and for condensates treated with RGD-functionalized dendrimers in solution (integrin blocking), after 6 days of chondrogenic induction (18 ≤ n ≤ 21). **b)** Schematics of the transplantation assay. Cell condensates formed on S_90_ and S_18_ substrates were collected at day 3 of chondrogenic induction and plated on fresh S_90_ or S_18_ substrates for 3 more days of chondrogenic induction. **c)** Quantification of the relative area immunostained for Cx43 for non-transplanted and transplanted condensates (12 ≤ n ≤ 17). Results are given as the mean ± SEM, **p < 0.01.

We then questioned whether variations in the substrate input would produce any changes on already formed cell condensates. Therefore, we designed a transplantation assay in which cell condensates on S_90_ and S_18_ were collected at day 3 of chondrogenic induction, plated on new S_90_ and S_18_ substrates and maintained for 3 more days in chondrogenic culture (Fig. 4b). The percentage of area immunostained for Cx43 increased by 71 ± 21% in cell condensates transplanted from S_90_ to fresh S_90_, and by 61 ± 19% in cell condensates transplanted from S_18_ to S_90_. Transplantation of either S_90_ or S_18_ cell condensates to fresh S_18_ substrates did not render significant changes in Cx43 immunostained area (Fig. 4c).

Cell condensates formed on one substrate and then transplanted to another will sense the new input locally, only from the side in direct contact with the new substrate, and only basal cells will receive it directly. Hence, we considered if transplantation effects are confined to the basal cell layer, or they can propagate inside the cell condensate up to the apical layer. Transversal views of the cell condensates transplanted to S_90_ nanopatterns showed that the increase in Cx43 staining is visible among different heights inside the cell condensate (Fig. 5a). Moreover, the proportion of Cx43 between the basal versus apical regions was equal in non-transplanted and transplanted cell condensates, indicating that transplantation did not alter the ratios of protein distribution among layers (Fig. 5b). As perturbations at the cell membrane are transduced into chemical responses by their propagation from integrins through the cytoskeleton [18], we hypothesized that propagation of the substrate input inside the cell condensate may involve actomyosin contraction [51,52]. To assess the effects of myosin-2 activity, we treated cell condensates of 6 days of chondrogenic induction on S_90_ and S_18_ nanopatterns with the myosin-2 inhibitor blebbistatin. Blebbistatin treatment led to a diffuse distribution of actin in cell condensates, instead of concentrating in clearly defined fibers as observed in control conditions (Fig. 5c), and significantly decreased the percentage of area immunostained for Cx43 in S_90_ cell condensates, while no changes were observed for S_18_ (Fig. 5d).

**Figure 5.**
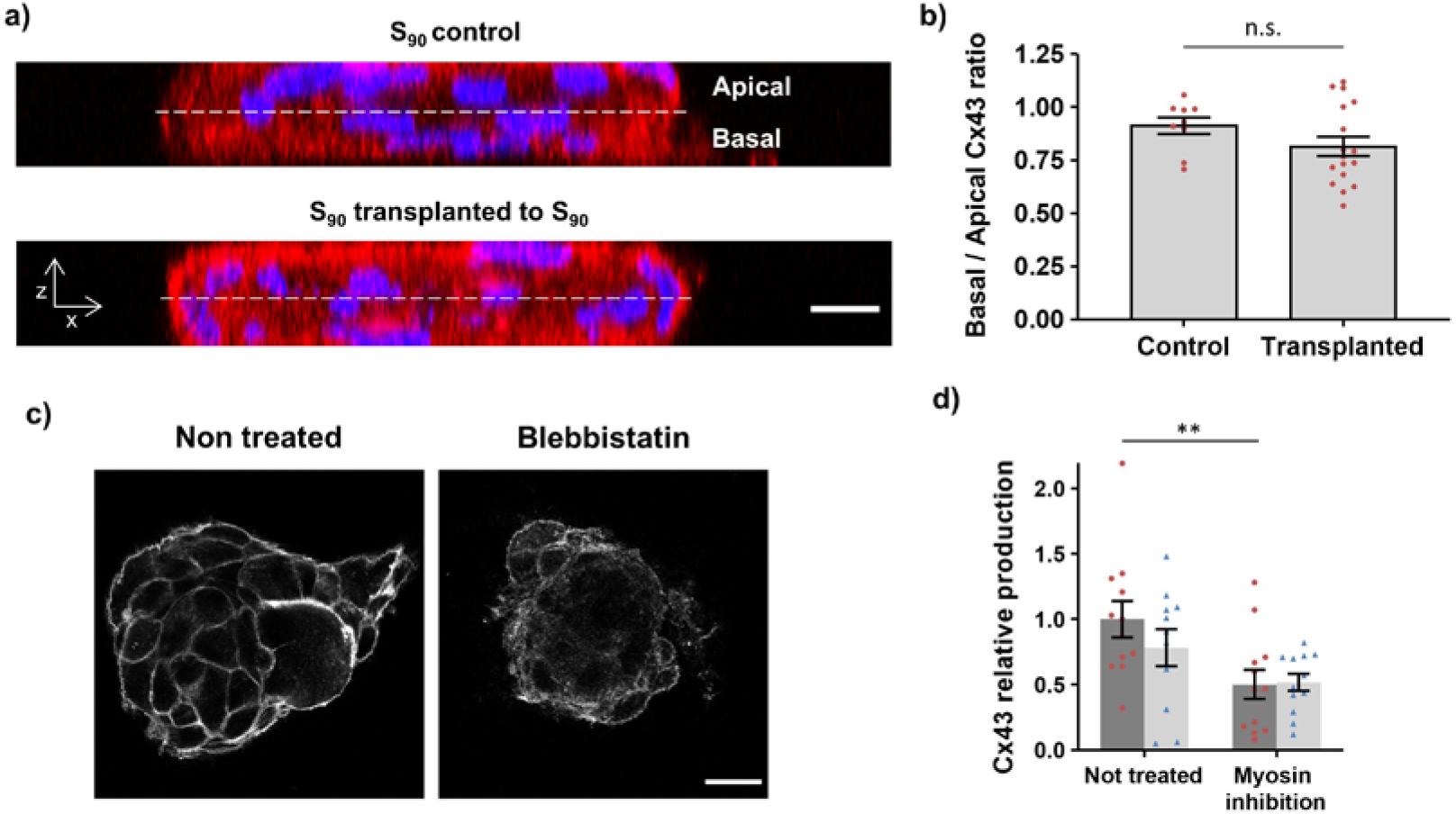
**a)** Confocal side views of control and transplanted S_90_ cell condensates, stained for Cx43 (red) and cell nuclei (Hoechst; blue). Scale bar = 10 μm. **b)** Ratio between the percentage of area immunostained for Cx43 at the basal and apical regions of S_90_ condensates in control substrates and in S_90_ cell condensates transplanted to fresh S_90_ nanopatterns (9 ≤ n ≤ 17). **c)** Actin staining for S_90_ cell condensates at day 6 of chondrogenic induction without (non-treated) and with blebbistatin (myosin inhibition). Scale bar = 25 μm. **d)** Quantification of the relative area immunostained for Cx43 on S_90_ (dark bars with red dots) and S_18_ (light bars with blue triangles) obtained for non-treated cell condensates and for condensates treated with blebbistatin, after 6 days of chondrogenic induction (n = 12). Results are given as mean ± SEM, **p < 0.01.

## Discussion

We have previously developed a technique to produce RGD-dendrimer-based nanopatterns with liquid-like order and controlled spacing on large surfaces areas, thus being fully compatible with normal cell culture protocols [31]. These nanopatterns allowed us to control cell adhesion and differentiation of hASCs towards different lineages [31–34,36]. We previously found that S_90_ nanopatterns produce larger and mechanically more stable prechondrogenic cell condensates [32,33,36]. Cells on pristine and nanopatterned substrates immediately start aggregating, while cells on FN coatings first proliferate and adopt a monolayer configuration, from which a few small cell condensates develop after 3 to 5 days in culture [32]. The size of the cell condensates increases with surface adhesiveness on the nanopatterns during the first 5 days of chondrogenic induction [32]. They maintain their three-dimensional (3D) structure in culture up to day 9, and then they start disaggregating. Only cell condensates on S_90_ retain their 3D structure up to day 14. The increase in mechanical stability of cell condensates on S_90_ nanopatterns was attributed to a balance between integrin-mediated cell cohesion, given by the nanopattern configuration, and the surface tension of the cell condensates (measured as N-cadherin expression) [36].

Here we further investigated cell-cell cohesion in the cell condensates produced on the different nanopattern configurations (S_18_, S_45_ and S_90_, from lower to higher adherence) and on the negative (S_0_) and positive (S_FN_) controls for cell adhesion. We observed increased cell compaction (proximity among the nuclei of adjacent cells) in cell condensates on high adherent substrates, S_90_ and S_FN_. Shorter intercellular distances in condensates at day 6 of chondrogenic induction could be a factor explaining the observed resistance of large S_90_ condensates to collapse through day 14. Moreover, the adherens junction protein VCL shows a tendency to increase with decreasing substrate adhesiveness. This agrees with the results from Byers and co-workers, who reported that VCL was essential to confer cell cohesiveness to embryonal carcinoma cell condensates obtained under non-adherent substrate conditions [53]. Many cell types, such as fibroblasts or epithelial cells, tend to aggregate when cultured on low-adherence or soft substrates, but migrate out of explants if cultured on stiff substrates [54]. Thus, the increasing the amount of VCL obtained with decreasing substrate adherence may be attributed to a response of cells to the lack of cell-substrate interactions, and as an alternative source of stability during cell condensation. This observation is supported by immunofluorescence images (Fig. 1d) showing that VCL was more present at the rim of cell condensates, bundling the cells in a compacted structure, similarly as observed in amnioserosa cells of embryos that undergo the late stages of dorsal closure, and in which VCL confers cell cohesiveness and maintains tension [55].

The process of mesenchymal cell condensation during chondrogenesis is concurrent with the formation of an extensive GJ network, which allows intercellular communication and synchronization during tissue patterning [5–9]. Since they bring the membranes of adjacent cells closer together, they also improve the mechanical stability at cell-cell junctions [56] and can act as cell-adhesive sites themselves [57–59]. Connective tissues, such as cartilage, particularly rely on this form of intercellular communication for successful development and homeostasis [8,9,60]. We measured the expression and production of Cx43, a connexin that is ubiquitously expressed in developing cartilage [11,61]. Results show that Cx43 was significantly higher on S_90_ cell condensates at day 6 of chondrogenic induction, although it tends to level off among the different nanopattern configurations and S_0_ at day 9. Cx43 facilitates the formation of multicellular aggregates [58] and mediates cell-cell adhesion during self-assembly of microtissues from human granulosa cells and fibroblasts [62], which could also contribute to the increased mechanical stability of S_90_ cell condensates.

Immunostaining of Cx43 showed that aggregates of different sizes (1-10 μm in diameter) are formed at different locations in cells within the mesenchymal cell condensates. We skeletonized Cx43 confocal images to visualize the 3D distribution of the protein forming clusters, either GJs or arrayed connexons. The analyses of EPVs showed that network connectivity increased with substrate adhesiveness up to S_90_ and then decreased for S_FN_, and that at day 9 of chondrogenic induction S_90_ condensates presented a more intricate network of assembled Cx43. To assess whether Cx43 was assembled into functional GJ plaques, we conducted a NB tracer assay, which resulted in a significantly higher uptake for S_90_ at day 6 of chondrogenic induction, with a calculated value of 7.7 μm/min. This was found within the range of previously reported values for HeLa cells, which go from 6 to 14 μm/min [42]. Given a condensate area of 10000 μm^2^ for S_90_, and hence an average radius of approximately 56 μm (if we consider a circular shape), the measured uptakes also explain why an exposure time of 10 minutes, at the calculated diffusion rate for the neurobiotin tracer of 7.7 μm/min, results in tracer-filled condensates, as observed. We also conducted FRAP measurements on S_90_ and S_18_ cell condensates. For S_90_, a 30% of fluorescence recovery was obtained after FRAP quantitative analysis, with a T half of 5.3 min, in agreement with previously reported results of gap fluorescence recovery after photobleaching [42,48]. To specifically address GJ-exchange, we conducted an analogous experiment with S_90_ cell condensates pre-treated with the GJ inhibitor 18βGA. We observed that FRAP was drastically reduced with values of fluorescence recovery similar to S_18_, thus confirming that the fast recovery in S_90_ can be attributed to GJ coupling and suggesting that in S_18_ cell condensates GJ coupling is very low or negligible, in line with the results obtained in the NB tracer assay.

Results show that cell condensates on S_90_ nanopatterns developed a more efficient Cx43-based GJ network, thus suggesting integrin-based adhesions play a role in the development of GJIC during mesenchymal cell condensation in chondrogenesis. Similarly, hMSCs from apical papilla respond to increasing substrate stiffness by assembling gap junction plaques, resulting in increased transmission of lucifer yellow tracer, through a process regulated by FAK and paxillin binding to Cx43 [15]. This would correspond with increased paxillin expression previously observed also on S_90_ substrates [32,36].

To directly assess the role of integrins, we conducted a competitive assay in which substrate (RGD)-integrin interaction was perturbed by the addition of RGD-functionalized dendrimers in solution. We analysed the percentage of area immunostained for Cx43 on cell condensates of S_90_ and S_18_ nanopatterns after 6 days of chondrogenic induction with and without RGD-functionalized dendrimers in solution. We found a significant decrease in the percentage of area immunostained for Cx43 only in S_90_. This observation, together with results of condensate stability and GJIC, indicates that the establishment of a functional GJ network during hASCs chondrogenic condensation is an adhesion-gated mechanism, in which integrin-mediated cell adhesion affects production and assembly of GJs. Cell response is triggered when integrins are engaged through the S_90_ nanopattern configuration. Thus, the S_90_ configuration provides the required adhesion threshold for cell response. This is also supported by our previous results showing that cell condensates on S_90_ nanopatterns are the only ones able to enter the early stages of chondrogenic commitment after 3 days of chondrogenic treatment [32,36].

We also conducted a transplantation assay to test whether changes in the substrate input will produce any changes on already formed cell condensates. Transplantation of cell condensates from S_18_ and S_90_ nanopatterns to a new S_90_ substrate caused a significant increase in Cx43, suggesting that cell condensates still preserve a certain level of mesenchymal plasticity at early stages of chondrogenic differentiation [63], thereby allowing phenotype reconfiguration in response to the new ECM input. The fact that transplantation from S_90_ to new S_90_ caused a further increase in Cx43 suggests that cells exert a certain degree of substrate remodelling that masks the original substrate information after at least three days of culture. This could be either due to matrix secretion, which would progressively cover the substrate and offer competing adhesion sites to the cells; or to changes in the density and distribution of nanopatterned ligands, given that RGD dendrimers are adsorbed but not covalently bound to the substrates. Upon transplantation to fresh substrates of the optimal ligand density (S_90_), cell condensates are exposed again to the original input, cell signalling is triggered, and they respond with further Cx43 production. No effects were observed for transplantation to S_18_ substrates, as expected.

We observed that the increase in the percentage of area immunostained for Cx43 can be noticed at any level inside the cell condensates, thus indicating that the effects of the substrate are not confined to basal cells, but they propagate inside the tissue. As integrin transduction implicates the cell cytoskeleton [18], we conducted a myosin-2 inhibition assay to see whether actomyosin contraction in the cell cortex may influence Cx43. Results showed that treatment with blebbistatin (a myosin-2 inhibitor) caused a decrease in the percentage of area immunostained for Cx43 in S_90_ cell condensates. This is in agreement with previous reports showing that connexin recruitment in gap junctions is modulated by their interactions with cytoskeletal structures [64,65]. We propose that once integrins are engaged, the adhesion information from the substrate is transduced and propagates through the actin filaments by myosin-2 mediated contraction, which in turn can regulate GJ accretion through ZO-1 [14]. Similarly, Cx43 accumulation into GJs and the subsequent establishment of GJIC has been reported to rely on myosin-VI in cell monolayers [66]. In human tenocytes, Cx43 co-localizes with actin only under myosin-II activity [67], showing that mechanical contractions are necessary for gap junctional regulation through cytoskeletal proteins. These results also agree with previous studies indicating that the dynamics of membrane proteins, such as connexins, are regulated by the cell’s actin cortex [68].

Overall, results show that integrin-based cell-matrix adhesions have an impact on the development of GJIC during mesenchymal cell condensation in chondrogenesis. In particular, we observed that the S_90_ nanopattern configuration produces a threshold signal for cell response enhancing mechanical stability and intercellular communication in mesenchymal cell condensates, which correlate with improved chondrogenic differentiation [32,33]. Clausen-Schaumann and co-workers have investigated in a mouse model the structural and mechanical properties of developing cartilage in the proliferative zone of the tibial growth plate, by atomic force microscopy (AFM) [69]. AFM topography images of samples from the ECM in between cells at E13.5 showed that it is organized in a randomly oriented meshwork of fibrils. Authors calculated a collagen fibril density of 8.8-10.5 fibrils/μm^2^ in these samples and they found it increases at later developmental stages. A distance between the fibrils of around 100 nm suggested this could be the maximum distance between two neighboring binding sites for integrins. The collagen fibrils contain integrin binding regions mostly located within the overlapping regions (axial D-periodicity of 67 nm) and the interfibrillar space could also contain other integrin-binding non-collagenous proteins [70,71]. Therefore, the S_90_ configuration, in which 90% of the surface area contains adhesion sites separated less than 70 nm, could be emulating the native disposition of integrin binding sites in the ECM of developing cartilage, thus favoring mesenchymal cell condensation and differentiation.

The results obtained have an immediate application to cartilage *in vitro* development [36,72], for which we are already producing cell constructs to be implanted in cartilage lesions of animal models, but they are also extensible to the study of other biological processes in which active ECM remodelling and thus, changes in the adhesion requirements and intercellular communication play an active role, such as in cancer progression [73].

## Conclusion

Our study shows that nanoscale ligand density regulates GJIC of developing cell condensations during chondrogenesis *in vitro*. A threshold nanopattern configuration in which the 90% of the surface area contains adhesion sites separated less than 70 nm activates cell signalling, enhancing both mechanical stability and GJIC. Similarities found in the nanospacing between integrin binding sites in native cartilage matrices, suggest this nanopattern configuration emulates the ECM adhesive requirements of developing cartilage, which could be applied to regenerative medicine purposes.

## Supporting information

permissions

## Acknowledgements

This work was supported by the Biomedical Research Networking Center (CIBER), Spain. CIBER is an initiative funded by the VI National R&D&i Plan 2008–2011, Iniciativa Ingenio 2010, Consolider Program, CIBER Actions, and the Instituto de Salud Carlos III (RD16/0006/0012; RD16/0011/0022), with the support of the European Regional Development Fund (ERDF). This work was funded by the CERCA Program and by the Commission for Universities and Research of the Department of Innovation, Universities, and Enterprise of the Generalitat de Catalunya (2017 SGR 1079). This work has been developed in the context of the AdvanceCat project (COMRDI15-1-0013) with the support of ACCIÓ (Catalonia Trade and Investment; Generalitat de Catalunya) under the Catalonian ERDF operational program 2014–2020. This work was funded by the Spanish Ministry of Economy and Competitiveness (MINECO) through the projects MINDS (Proyectos I+D Excelencia + FEDER): TEC2015-70104-P-P and BIOBOT (Programa Explora Ciencia / Tecnología): TEC2015-72718-EXP, as well as the Spanish Ministry of Science and Education (PID2019-104293GB-I00), the Spanish Ministry of Science and Innovation (Proyectos de I+D+I «Programación Conjunta Internacional», EuroNanoMed 2019 (PCI2019-111825-2), and the Consejería de Salud, Junta de Andalucía (UMA18-FEDERJA-007; UMA18-FEDERJA-133) and Consejeria de Transformacion Economica, Industria, Conocimiento y Universidades, Junta de Andalucia (PY20_00384). The authors further acknowledge funds from the INTERREG V cooperation program for Spain–Portugal (POCTEP) 2014–2020 project (0245_IBEROS_1_E). I. C. acknowledges support from MINECO through the Subsidies for Predoctoral Contracts for the Training of Doctors open call, co-funded by the European Social Fund (2016), grant number: BES-2016-076682. C. R-P. acknowledges support from i-PFIS Doctoral Program (IFI15/00151) funded by ISCIII. We thank C. Alcon for help in Western Blots, N. Montserrat for help with the transplantation assay, R. Paoli for producing Video S1, and P. Roca-Cusachs for the fruitful discussions. We also thank Dr. Bardia and Dr. Giakoumakis at the Advanced Digital Microscopy unit at Institute for Research in Biomedicine (IRB) for their guidance with the FRAP experiments and analysis.

