## Supplementary Information for "Nanoscale ligand density modulates gap junction intercellular communication of cell condensates during chondrogenesis"

### Nanoscale cell-matrix adhesions modulate gap junction intercellular communication of mesenchymal cell condensates during chondrogenesis (Supplementary Files)

Ignasi Casanellas[1,2,3], Anna Lagunas[3,1]*, Yolanda Vida[4,5], Ezequiel Pérez-Inestrosa[4,5], Cristina Rodríguez-Pereira[6], Joana Magalhaes[3,6], José A. Andrades[7,3,5], José Becerra[7,3,5], Josep Samitier[1,2,3]

[1]Institute for Bioengineering of Catalonia (IBEC), Barcelona Institute of Science and Technology (BIST). c/ Baldiri Reixac, 10-12, 08028 Barcelona, Spain.
[2]Department of Electronics and Biomedical Engineering, University of Barcelona (UB), Faculty of Physics. c/ Martí i Franquès, 1, 08028 Barcelona, Spain.
[3]Biomedical Research Networking Center in Bioengineering, Biomaterials, and Nanomedicine (CIBER-BBN). Av. Monforte de Lemos, 3-5. Pabellón 11. Planta 0, 28029 Madrid, Spain.
[4]Universidad de Málaga - IBIMA, Dpto. Química Orgánica. Campus de Teatinos s/n, 29071 Málaga, Spain.
[5]Centro Andaluz de Nanomedicina y Biotecnología-BIONAND. Parque Tecnológico de Andalucía, c/ Severo Ochoa 35, 29590 Campanillas, Málaga, Spain.
[6]Unidad de Medicina Regenerativa, Grupo de Investigación en Reumatología (GIR), Instituto de Investigación Biomédica de A Coruña (INIBIC), Complexo Hospitalario Universitario de A Coruña (CHUAC), Sergas, Universidade da Coruña (UDC). c/ Xubias de Arriba, 84, 15006 A Coruña, Spain.
[7]Department of Cell Biology, Genetics and Physiology, Universidad de Málaga (UMA), Instituto de Investigación Biomédica de Málaga (IBIMA). Av. Cervantes, 2, 29071 Málaga, Spain.

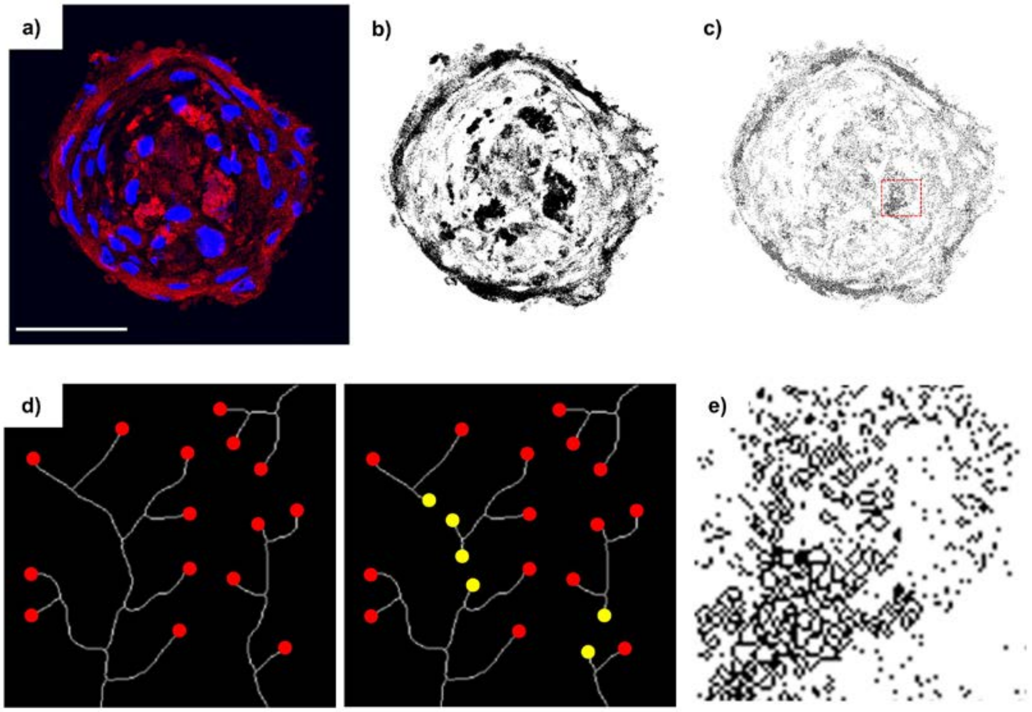


**Fig. S1: Analysis of Cx43 expression and intercellular communication network architecture**. **a)** Representative chondrogenic cell condensate stained for Cx43 (red) and cell nuclei (blue). **b)** Threshold selecting Cx43 expression.

**c)** Skeletonizing of the Cx43 network, from which end-point voxels and mean branch length were quantified. **a-c)** Scale bar = 50 µm. **d)** Schematic of network connectivity quantification as the inverse of the number of end-point voxels. White lines represent theoretical networks, red dots indicate end-point voxels, yellow dots indicate new end-point voxels as a results of network splitting. The network on the left is more extensively connected than the one on the right, and thus contains fewer end-point voxels. **e)** Zoomed-in section from the red square in **c**.

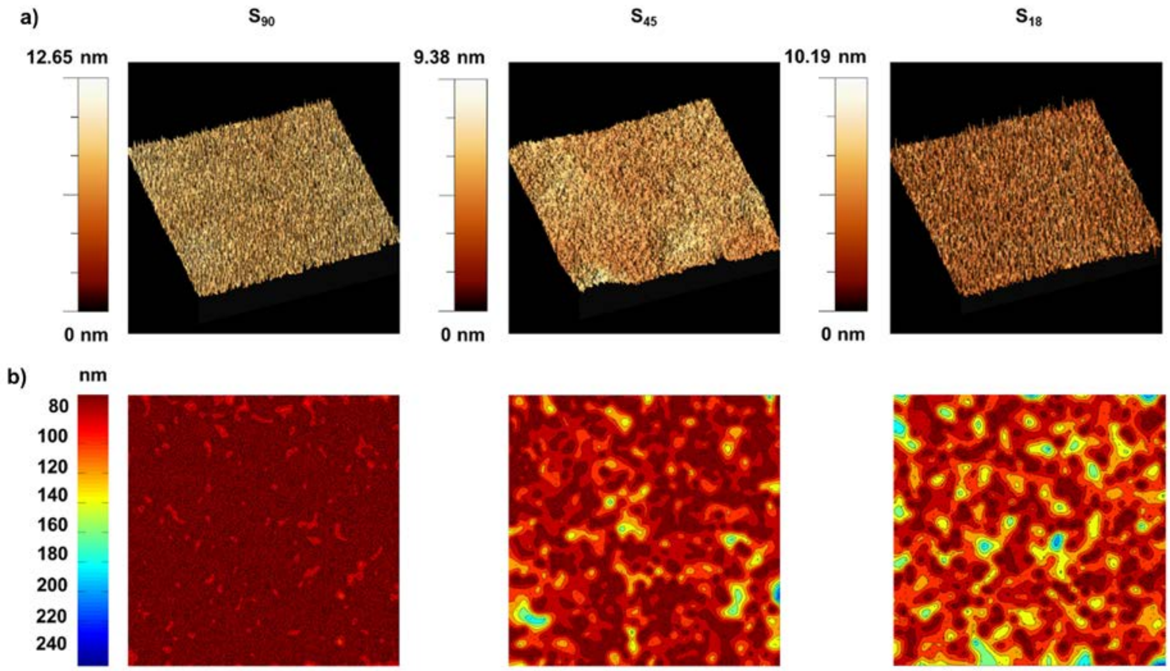


**Fig. S2: Substrates of tunable local surface adhesiveness obtained by nanopatterning RGD-functionalized dendrimers on PLLA**. **a)** Representative 3D AFM images (5x5 µm) of the nanopatterns' topography. **b)** Corresponding $d_{min}$ probability contour plots showing high-density RGD regions ($d_{min}$ < 70 nm) in dark red (dendrimer positions are superimposed in black for clarity). Adapted with permission from ref.30. Copyright (2017), Springer Nature: Springer Nature, Nano Research.

**Table S1: Nanopatterns adherent area**

| Nanopatterned substrate | Dendrimer concentration [% w w$^{-1}$] | % Adherent area ($d_{min}$ < 70 nm) |
|---|---|---|
| $S_{90}$ | 2.5 x $10^{-8}$ | 90 ± 2 |
| $S_{45}$ | $10^{-8}$ | 45 ± 7 |
| $S_{18}$ | 4 x $10^{-9}$ | 18 ± 11 |

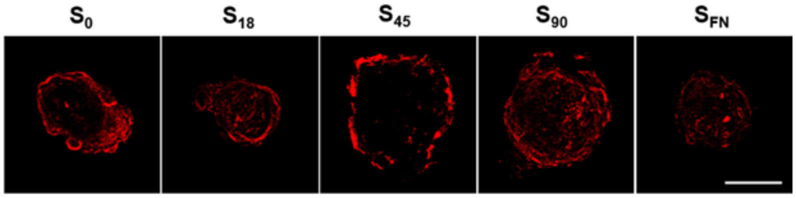


**Fig. S3:** Representative confocal z-projections of Cx43 immunostaining at day 6, scale bar = 40 µm.

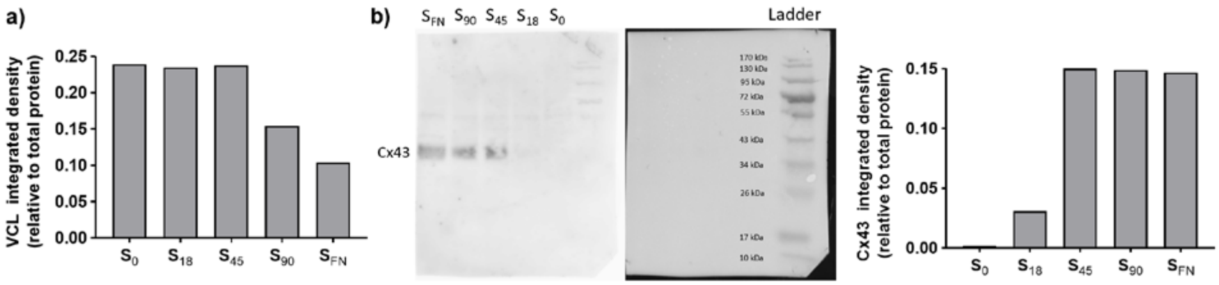


**Fig. S4: a)** Western blot quantification for vinculin at day 6 of chondrogenesis. **b)** Western blot and corresponding quantification for Cx43 at day 6 of chondrogenesis.

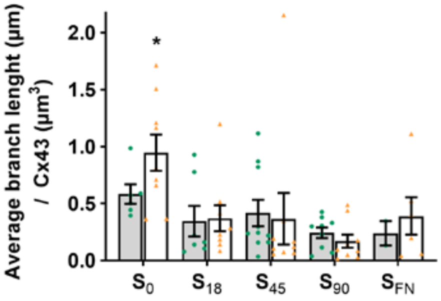


**Fig. S5:** Average length of Cx43 network branches in chondrogenic cell condensates (n≥2). Gray and white bars correspond to 6 and 9 days of chondrogenic induction, respectively. Green dots and orange triangles correspond to individual sample values at day 6 and day 9, respectively. Results given as the mean ± SEM., *$p < 0.05$

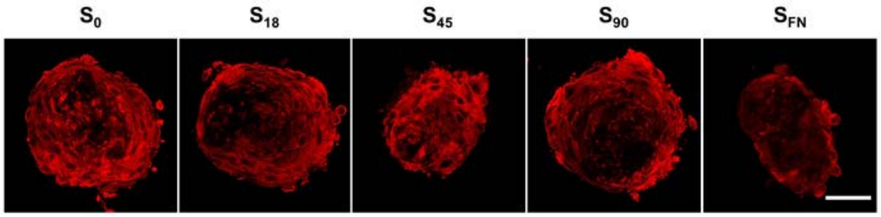


**Fig. S6:** Confocal z-projections showing neurobiotin tracer (red) after 10 min of exposure at day 6 of chondrogenic induction**.** Scale bar = 80 µm.
