## Supplementary material for "Nanoscale ligand density modulates gap junction intercellular communication of cell condensates during chondrogenesis": permissions

SPRINGER NATURE LICENSE
TERMS AND CONDITIONS

Jul 07, 2020

---

This Agreement between Anna Lagunas ("You") and Springer Nature ("Springer Nature") consists of your license details and the terms and conditions provided by Springer Nature and Copyright Clearance Center.

| | |
|---|---|
| License Number | 4863700739650 |
| License date | Jul 07, 2020 |
| Licensed Content Publisher | Springer Nature |
| Licensed Content Publication | Nano Research |
| Licensed Content Title | Tailoring RGD local surface density at the nanoscale toward adult stem cell chondrogenic commitment |
| Licensed Content Author | Anna Lagunas et al |
| Licensed Content Date | Dec 29, 2016 |
| Type of Use | Journal/Magazine |
| Requestor type | academic/university or research institute |
| Is this reuse sponsored by or associated with a pharmaceutical or a medical products company? | no |
| Format | electronic |

| | |
|---|---|
| Portion | figures/tables/illustrations |
| Number of figures/tables/illustrations | 1 |
| Will you be translating? | no |
| Circulation/distribution | 50000 or greater |
| Author of this Springer Nature content | yes |
| Title of new article | Nanoscale basal ligand density modulates gap junction intercellular communication network architecture during mesenchymal cell condensation |
| Lead author | Ignasi Casanellas |
| Title of targeted journal | ACS Applied Materials & Interfaces |
| Publisher | ACS Publications |
| Expected publication date | Sep 2020 |
| Portions | Figure 1b. dmin probability contour plot for 4 × 10–9% |
| Requestor Location | Anna Lagunas<br>Baldiri Reixac 10-12<br>Nanobioengineering Lab (IBEC).<br><br>Barcelona, 08028<br>Spain<br>Attn: Anna Lagunas |
| Customer VAT ID | ESG64045719 |
| Total | 0.00 EUR |

Terms and Conditions

**Springer Nature Customer Service Centre GmbH**
**Terms and Conditions**

This agreement sets out the terms and conditions of the licence (the **Licence**) between you and **Springer Nature Customer Service Centre GmbH** (the **Licensor**). By clicking 'accept' and completing the transaction for the material (**Licensed Material**), you also confirm your acceptance of these terms and conditions.

**1. Grant of License**

**1. 1.** The Licensor grants you a personal, non-exclusive, non-transferable, world-wide licence to reproduce the Licensed Material for the purpose specified in your order only. Licences are granted for the specific use requested in the order and for no other use, subject to the conditions below.

**1. 2.** The Licensor warrants that it has, to the best of its knowledge, the rights to license reuse of the Licensed Material. However, you should ensure that the material you are requesting is original to the Licensor and does not carry the copyright of another entity (as credited in the published version).

**1. 3.** If the credit line on any part of the material you have requested indicates that it was reprinted or adapted with permission from another source, then you should also seek permission from that source to reuse the material.

**2. Scope of Licence**

**2. 1.** You may only use the Licensed Content in the manner and to the extent permitted by these Ts&Cs and any applicable laws.

**2. 2.** A separate licence may be required for any additional use of the Licensed Material, e.g. where a licence has been purchased for print only use, separate permission must be obtained for electronic re-use. Similarly, a licence is only valid in the language selected and does not apply for editions in other languages unless additional translation rights have been granted separately in the licence. Any content owned by third parties are expressly excluded from the licence.

**2. 3.** Similarly, rights for additional components such as custom editions and derivatives require additional permission and may be subject to an additional fee. Please apply to for these rights.

**2. 4.** Where permission has been granted **free of charge** for material in print, permission may also be granted for any electronic version of that work, provided that the material is incidental to your work as a whole and that the electronic version is essentially equivalent to, or substitutes for, the print version.

**2. 5.** An alternative scope of licence may apply to signatories of the STM Permissions

Guidelines, as amended from time to time.

### 3. Duration of Licence

**3. 1.** A licence for is valid from the date of purchase ('Licence Date') at the end of the relevant period in the below table:

| Scope of Licence | Duration of Licence |
|---|---|
| Post on a website | 12 months |
| Presentations | 12 months |
| Books and journals | Lifetime of the edition in the language purchased |

### 4. Acknowledgement

**4. 1.** The Licensor's permission must be acknowledged next to the Licenced Material in print. In electronic form, this acknowledgement must be visible at the same time as the figures/tables/illustrations or abstract, and must be hyperlinked to the journal/book's homepage. Our required acknowledgement format is in the Appendix below.

### 5. Restrictions on use

**5. 1.** Use of the Licensed Material may be permitted for incidental promotional use and minor editing privileges e.g. minor adaptations of single figures, changes of format, colour and/or style where the adaptation is credited as set out in Appendix 1 below. Any other changes including but not limited to, cropping, adapting, omitting material that affect the meaning, intention or moral rights of the author are strictly prohibited.

**5. 2.** You must not use any Licensed Material as part of any design or trademark.

**5. 3.** Licensed Material may be used in Open Access Publications (OAP) before publication by Springer Nature, but any Licensed Material must be removed from OAP sites prior to final publication.

### 6. Ownership of Rights

**6. 1.** Licensed Material remains the property of either Licensor or the relevant third party and any rights not explicitly granted herein are expressly reserved.

### 7. Warranty

IN NO EVENT SHALL LICENSOR BE LIABLE TO YOU OR ANY OTHER PARTY OR

ANY OTHER PERSON OR FOR ANY SPECIAL, CONSEQUENTIAL, INCIDENTAL OR INDIRECT DAMAGES, HOWEVER CAUSED, ARISING OUT OF OR IN CONNECTION WITH THE DOWNLOADING, VIEWING OR USE OF THE MATERIALS REGARDLESS OF THE FORM OF ACTION, WHETHER FOR BREACH OF CONTRACT, BREACH OF WARRANTY, TORT, NEGLIGENCE, INFRINGEMENT OR OTHERWISE (INCLUDING, WITHOUT LIMITATION, DAMAGES BASED ON LOSS OF PROFITS, DATA, FILES, USE, BUSINESS OPPORTUNITY OR CLAIMS OF THIRD PARTIES), AND
WHETHER OR NOT THE PARTY HAS BEEN ADVISED OF THE POSSIBILITY OF SUCH DAMAGES. THIS LIMITATION SHALL APPLY NOTWITHSTANDING ANY FAILURE OF ESSENTIAL PURPOSE OF ANY LIMITED REMEDY PROVIDED HEREIN.

**8. Limitations**

**8. 1.** ***BOOKS ONLY:***Where **'reuse in a dissertation/thesis'** has been selected the following terms apply: Print rights of the final author's accepted manuscript (for clarity, NOT the published version) for up to 100 copies, electronic rights for use only on a personal website or institutional repository as defined by the Sherpa guideline (www.sherpa.ac.uk/romeo/).

**9. Termination and Cancellation**

**9. 1.** Licences will expire after the period shown in Clause 3 (above).

**9. 2.** Licensee reserves the right to terminate the Licence in the event that payment is not received in full or if there has been a breach of this agreement by you.

**Appendix 1 — Acknowledgements:**

**For Journal Content:**
Reprinted by permission from [**the Licensor**]: [**Journal Publisher** (e.g. Nature/Springer/Palgrave)] [**JOURNAL NAME**] [**REFERENCE CITATION** (Article name, Author(s) Name), [**COPYRIGHT**] (year of publication)

For **Advance Online Publication papers:**
Reprinted by permission from [**the Licensor**]: [**Journal Publisher** (e.g. Nature/Springer/Palgrave)] [**JOURNAL NAME**] [**REFERENCE CITATION** (Article name, Author(s) Name), [**COPYRIGHT**] (year of publication), advance online publication, day month year (doi: 10.1038/sj.[JOURNAL ACRONYM].)

**For Adaptations/Translations:**
Adapted/Translated by permission from [**the Licensor**]: [**Journal Publisher** (e.g. Nature/Springer/Palgrave)] [**JOURNAL NAME**] [**REFERENCE CITATION** (Article name, Author(s) Name), [**COPYRIGHT**] (year of publication)

**Note: For any republication from the British Journal of Cancer, the following credit line style applies:**

Reprinted/adapted/translated by permission from [**the Licensor**]: on behalf of Cancer Research UK: : [**Journal Publisher** (e.g. Nature/Springer/Palgrave)] [**JOURNAL NAME**] [**REFERENCE CITATION** (Article name, Author(s) Name), [**COPYRIGHT**] (year of publication)

For **Advance Online Publication** papers:
Reprinted by permission from The [**the Licensor**]: on behalf of Cancer Research UK: [**Journal Publisher** (e.g. Nature/Springer/Palgrave)] [**JOURNAL NAME**] [**REFERENCE CITATION** (Article name, Author(s) Name), [**COPYRIGHT**] (year of publication), advance online publication, day month year (doi: 10.1038/sj. [JOURNAL ACRONYM])

**For Book content:**
Reprinted/adapted by permission from [**the Licensor**]: [**Book Publisher** (e.g. Palgrave Macmillan, Springer etc) [**Book Title**] by [**Book author**(s)] [**COPYRIGHT**] (year of publication)

**Other Conditions**:

Version 1.2

**Questions? or +1-855-239-3415 (toll free in the US) or +1-978-646-2777.**
